# Effects of artificially introduced *Enterococcus faecalis* strains in experimental necrotizing enterocolitis

**DOI:** 10.1101/623512

**Authors:** Patrick T. Delaplain, Brandon A. Bell, Jin Wang, Mubina Isani, Emily Zhang, Christopher P. Gayer, Anatoly V. Grishin, Henri R. Ford

## Abstract

*Enterococcus faecalis* is a ubiquitous intestinal symbiont and common early colonizer of the neonatal gut. Although colonization with *E. faecalis* has been previously associated with decreased NEC pathology, these bacteria have been also implicated as opportunistic pathogens. Here we characterized 21 strains of *E. faecalis*, naturally occurring in 4-day-old rats, for potentially pathogenic properties and ability to colonize the neonatal gut. The strains differed in hemolysis, gelatin liquefaction, antibiotic resistance, biofilm formation, and ability to activate the pro-inflammatory transcription factor NF-κB in cultured enterocytes. Only 3 strains appreciably colonized the neonatal intestine on day 4 after artificial introduction with the first feeding. The best colonizer, strain BB70, effectively displaced maternal *E. faecalis* and significantly increased NEC pathology. Our results show that colonization with *E. faecalis* may predispose neonates to NEC.

## Introduction

Necrotizing enterocolitis (NEC) affects approximately 1 in 1000 live births and is one of the leading causes of mortality among preterm infants [1]. Although the pathogenesis of this disease is not yet fully understood, it is broadly accepted that bacterial colonization of the immature intestine in combination with perinatal stresses such as formula feeding, hypoxia, and hypothermia play the leading role [2–4]. While no single pathogen is likely responsible for NEC, previous work has implicated clostridia [5], *Cronobacter* [6, 7], *E. coli* [8], and lactobacilli [9–11] as either NEC-promoting or protective colonizers of the neonatal intestine. Importantly, protective or pathogenic properties of these bacteria are strain-specific.

Probiotics, bacteria believed to be beneficial upon administration, have been extensively tried for prevention of NEC. Most of these trials are encouraging [12–14]. However, probiotics may cause adverse effects [15]. Evidence-based recommendations for clinical use of probiotics in NEC have not yet been developed due to lack of standardization of bacterial species/strains, doses, and treatment regimens across different trials [16]. A rational approach to probiotic therapy would be to identify commensals that effectively colonize the neonatal gut upon introduction and protect from NEC in animal models.

*E. faecalis* is a bacterial species of potential relevance to NEC. These bacteria constitute up to 1% of adult intestinal microbiome [17] and are readily transmitted from mothers to neonates in both humans [18, 19] and rodents [20, 21]. NEC patients tended to harbor lower percentages of *E. faecalis* in their microbiomes compared to healthy controls, but this tendency was not significant [22, 23]. Importantly, *E. faecalis* has been also implicated as pathogen [24]. To gain insight into potential role of *E. faecalis* in the pathogenesis of experimental NEC, we isolated multiple strains of these bacteria from 4-day old rats and examined their ability to colonize the neonatal intestine and to alter NEC pathology. Only few strains colonized the intestine following artificial introduction with first feeding. The best colonizing strain significantly exacerbated NEC pathology.

## Materials and Methods

### NEC Model

All animal experiments were approved by the CHLA Institutional Animal Care and Use Committee (IACUC). Timed-pregnant Sprague Dawley rats were obtained from either Envigo (Placentia, CA) or Charles River Laboratories (Hollister, CA). Newborn rats were separated from dams at birth and were kept in an infant incubator (Ohio Medical Products, Madison, WI) at 30°C and 90% humidity. NEC was induced by formula feeding and hypoxia, according to our previously published protocol [20, 25, 26]. Neonatal rats are fed 200 μl of formula (15 g Similac 60/40, Ross Pediatrics Columbus, OH in 75 ml of Esbilac canine milk replacement, Pet-Ag Inc., Hampshire, IL) every 8 h for 4 d. Fresh formula is prepared daily, each new batch is tested for bacterial contamination by plating on blood agar and MRS, and care is taken with each feeding to prevent introduction of extraneous bacteria. Pups are subjected to hypoxia at the conclusion of each feeding (10 min in 95% N_2_ and 5% O_2_). On day 4, animals are euthanized by decapitation and terminal ileum is collected for NEC pathology score and plating of intestinal contents. Samples for pathology scoring are fixed in formalin, embedded in paraffin, sectioned and stained with hematoxylin-eosine. These are then scored by a pathologist blinded to treatment groups. NEC score is assigned based on the degree of observed injury to the intestinal epithelium based on a 5-point scale (0: no pathology; 1: epithelial sloughing and/or mild sub-mucosal edema; 2: damage to the tips of the villi; 3: damage to more than half of the villi; 4: complete obliteration of the epithelium). Samples collected for bacterial analysis are homogenized in PBS, serially diluted and plated onto diagnostic media within 2 h of collection. Adult animals are euthanized by CO_2_ asphyxiation. If animals were treated with *E. faecalis*, bacteria were resuspended in formula and given with the first feed.

### Identification of bacteria

Independent isolates of *E. faecalis* were established from the intestinal contents of 4-day-old rats, colony-purified and kept as frozen stock as described previously [20]. To characterize populations of intestinal bacteria, freshly excised intestines were homogenized, serially diluted, and plated on blood agar (Sigma) for broad range of bacteria and MRS agar (Oxoid, Basinstoke, UK) for lactic bacteria. Plates were incubated for 4 d at 37°C in air (blood agar) or CO_2_ atmosphere (MRS agar). Emerging colonies were classified according to their appearance. Pure cultures for each colony type were purified by re-streaking and kept as frozen stocks. Bacterial species were identified by sequencing 16S rRNA gene PCR-amplified with 27F and 1492R primers at GeneWiz (Los Angeles, CA). Sequences were queried against NCBI non-redundant nucleotide (nt) database using the BLAST algorithm.

### Bacterial culture and phenotypes

*E. faecalis* bacteria were grown at 37°C aerobically in Brain Heart Infusion (BHI), Tryptic Soy Broth (TSB), or Luria Broth (LB). For pouring plates, agar was added to 17 g/L. Selective media for isolating *E. faecalis* contained 0.4 g/L sodium azide. *E. faecalis* phenotypes were determined by replica plating onto diagnostic media including blood agar, gelatin liquefaction medium (5 g/L peptone, 3 g/L beef extract, 120 g/L gelatin), antibiotic agar (LB supplemented with 50 mg/L ampicillin, or 100 mg/ml kanamycin, or 30 mg/l rifaximin), β-galactosidase agar (LB supplemented with 30 mg/L X-gal and 2 mM IPTG), and sugar fermentation agar (LB supplemented with 0.2 mg/L Neutral Red and 1% appropriate sugar). Gelatin plates were incubated upright at room temperature. Bacterial culture density was determined by spectrophotometry at 600 nm. Correlation between OD_600_ and CFU/ml was determined by serial dilution and plating.

### Restriction endonuclease analysis of bacterial DNA

Bacterial DNA was extracted from overnight culture by 5-min vortexing with 200 μm glass beads in TEN buffer (50 mM Tris pH 8.0, 100 mM NaCl, 10mM EDTA), overnight digestion at 50°C following addition of SDS and Proteinase K to 1% and 20 μg/ml, respectively, phenol/chloroform extraction, and ethanol precipitation. 5 μg DNA samples were digested with 10 u *Hind*III (New England Biolabs, Ipswich, MA) for 2 h at 37°C. Digestion products were resolved by electrophoresis through 0.8% agarose Tris-acetate gel. Gels were stained with ethidium bromide and photographed under UV light using GelDoc XR (Bio-Rad, Hercules, CA).

### Biofilm formation

A modified crystal violet assay, as previously described [27–29], was used to quantify biofilm formation. Overnight cultures of *E. faecalis* were diluted 1:50 in fresh medium and inoculated into wells of a 96-well polystyrene plate. Following static 24 h incubation at 37°C, plate was rinsed 3 times with PBS and air dried. After 10 min fixing with 3:1 ethanol:acetic acid, biofilms were stained with 0.1% crystal violet for 15 min. Wells were then washed with water until effluent ran clear. Crystal violet was extracted with 10% acetic acid, samples transferred to a new 96-well plate and OD_550_ was measured on plate reader.

### Binding of bacteria to enterocytes and activation of NF-κB

IEC-6 cells (rat intestinal epithelial cells) were obtained from ATCC and grown in DMEM+5% FBS as recommended by the supplier. Cells (passage 21-30) were used at 70-90% confluence. Bacteria grown overnight were diluted in DMEM and added to IEC-6 cells. After 30 min incubation at 37°C, cells were rinsed 3 times with ice-cold PBS, collected, serially diluted and plated on blood agar for bacterial quantification.

For activation of NF-κB, IEC-6 cells were treated with bacteria for 15 min, lysed on ice for 10 min with RIPA buffer (50 mM Tris-HCl pH 7.0, 100 mM NaCl, 1% NP-40, 0.5% sodium deoxycholate, 0.1% SDS, 1 mM PMSF). Lysates cleared by centrifugation at 10,000g for 10 min were mixed with 2x Laemmli buffer and boiled for 1 min. 20 μg protein samples were resolved on a 10% SDS-polyacrylamide gel. Transfer of protein onto nitrocellulose membrane, membrane blocking, incubation with primary antibody for IκBα (Cell Signaling, Danvers, MA) and secondary HRP-conjugated antibody were performed as recommended by antibody supplier. After extensive washing in PBS, membranes were impregnated with peroxide-luminol reagent and exposed to x-ray film.

### Statistics

Means for parametric data were compared using unpaired 2-sample *t*-test. Categorical and ordinal data were compared using χ^2^ test. All analyses were conducted in *R* v3.5.1 [30]. Graphics were designed either in R or GraphPad Prism v8.

## Results

### Diversity of *Enterococcus faecalis* in 4-day old rats

To examine potential relationship between *E. faecalis* and NEC, we sought to isolate enterococci naturally occurring in rats, identify different strains, and examine their properties in the rat model of NEC. Enterococci were isolated by blood agar plating of intestinal content from 4-day-old rats subjected to the NEC-inducing regimen of formula feeding and hypoxia. *E. faecalis* were identified by their resistance to azide, characteristic morphology upon Gram staining, and 16S rRNA gene sequencing. In our previous study, *Enterococcus* spp. was found in about 90% of intestinal samples. In the animals where enterococci were found, they constituted 17±3% of the bacterial populations [20]. To characterize diversity of *E. faecalis*, we examined 147 independent isolates of these bacteria collected from 4-day-old offspring of Charles River and Envigo (formerly Harlan) rats during 2008-2013. All isolates were catalase-negative, glucose-, fructose-, and sucrose-fermenting, ampicillin- and tetracycline-sensitive. Characteristics that differed among isolates included colony morphology, hemolysis, gelatin liquefaction, β-galactosidase activity, resistance to kanamycin or rifaximin, and fermentation of sorbitol, mannitol, and arabinose (**S1 Data File, Fig 1A**). Each unique combination of these phenotypes defined a distinct strain. We thus identified 21 different strains of *E. faecalis*, each represented by between 1 and 59 isolates (**S2 Data File, Fig 1B**). All strains belonged to one of the two genomic groups, as revealed by patterns of Hin*d*III DNA fragments (**S2 Data File, Fig 1C**). These results indicate that despite being kept at specific pathogen-free environment at facilities renowned for their high standards of care, laboratory rats harbor and transfer to their offspring a diverse array of *E. faecalis* strains.

**Fig 1.**
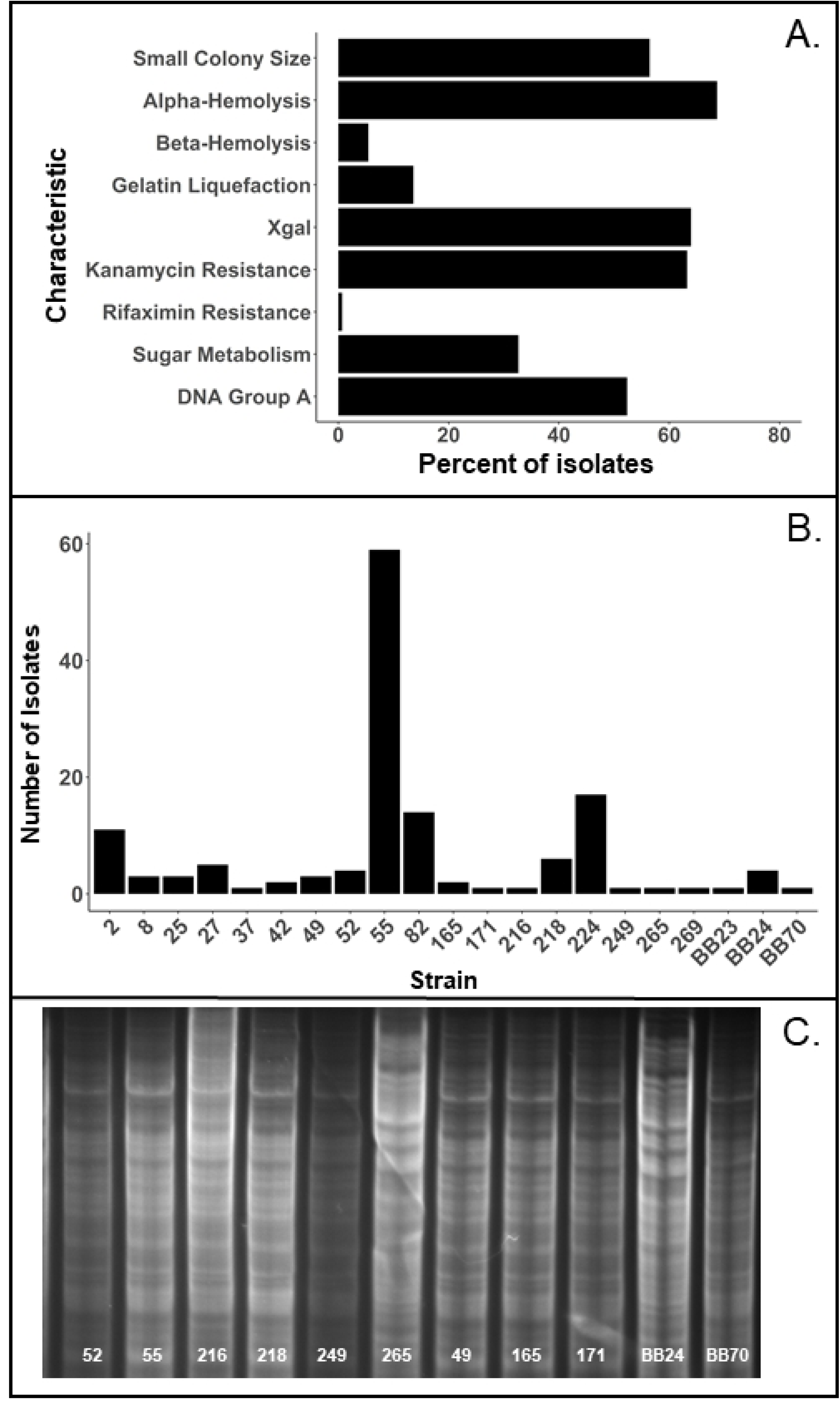
Diversity of *E. faecalis* in rats. Frequencies of different phenotypes (**A**) and different strains (**B**) within a group of *E. faecalis* isolates (n = 147) from 4-day old rats. (**C**) Patterns of genomic DNA Hin*d*III fragments of indicated strains. Note dissimilarity of DNA patterns A (55-249, 49-171, and BB70) and B (265, BB24).

### Potential pathogenic properties of *E. faecalis* strains identified in vitro

In order to narrow down the list of *E. faecalis* strains for in vivo studies, we examined strains’ potential pathogenic properties. Of the phenotypes described above, antibiotic resistance [31], hemolysis, and proteolysis [32] may contribute to pathogenicity. Another pathogenic phenotype of relevance to NEC could be the ability of bacteria to trigger mucosal inflammatory response. To characterize this phenotype, we treated IEC-6 cells (intestinal epithelioid cell line of rat origin) with each of the 21 strains of *E. faecalis* and examined activation of the pro-inflammatory transcription factor NF-κB by western blotting for the inhibitory subunit IκBα. Strains 25, 37, 49, and 82 caused degradation of IκBα (i.e. activation of NF-κB), whereas other strains caused partial degradation or no degradation (**Fig 2**).

**Fig 2.**
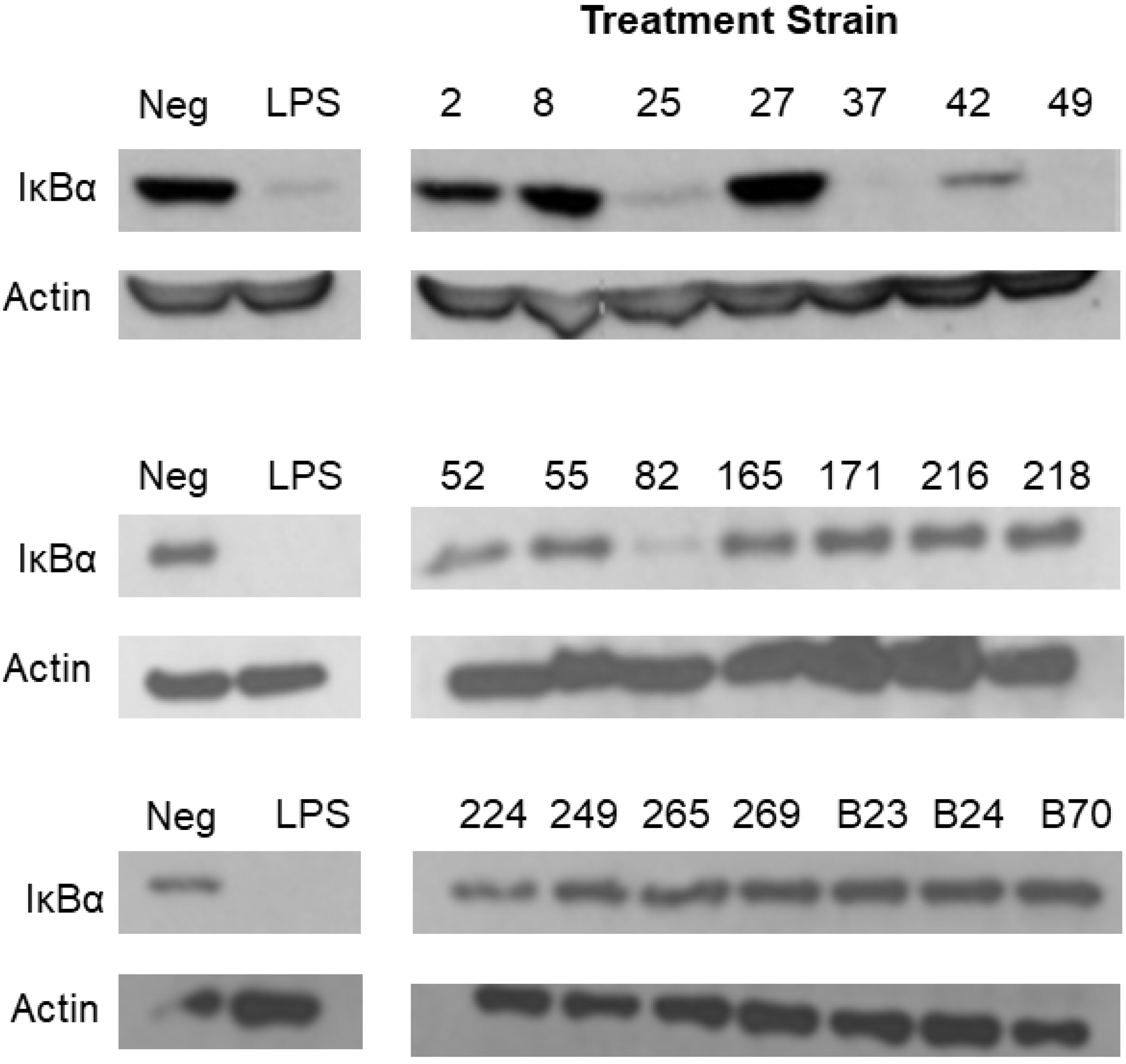
Activation of NF-κB by different strains of *E. faecalis*. IEC-6 cells were treated with each strain of *E. faecalis* and activation of the NF-κB pathway was determined by western blotting for IκBα. β-actin reprobes are included to demonstrate lane load. Representative blots of 3 independent experiments are shown.

Efficient binding to target cells may be one more phenotype associated with pathogenicity [7]. To characterize binding of our *E. faecalis* strains to enterocytes, IEC-6 monolayers were incubated with bacteria, washed, and homogenized. Resulting homogenates were serially diluted and plated onto BHI-azide agar for *E. faecalis* counts. There were no significant differences in binding efficiency of different strains (data not shown). On average, 11±1.4% of 10^8^ CFU input, or 0.32±0.04 CFU per IEC-6 cell were bound. Binding was weak – numbers of bound bacteria progressively decreased with additional washes (data not shown).

Biofilm formation, which may be associated with efficient colonization [21], is yet another potentially pathogenic phenotype. We measured biofilm formation in our *E. faecalis* strains by overnight culturing in polystyrene plates, washing off suspended bacteria, and biofilm staining with crystal violet (**Fig 3A**). Biofilm formation varied considerably among strains and was not associated with other phenotypes. Although BHI is a recommended culture medium for *E. faecalis*, it promoted the lowest average biofilm formation across strains compared to TSB or LB (**Fig 3B**). Thus, our strains of *E. faecalis* differed in inflammatory activation and biofilm formation properties, but not in enterocyte binding efficiency.

**Fig 3.**
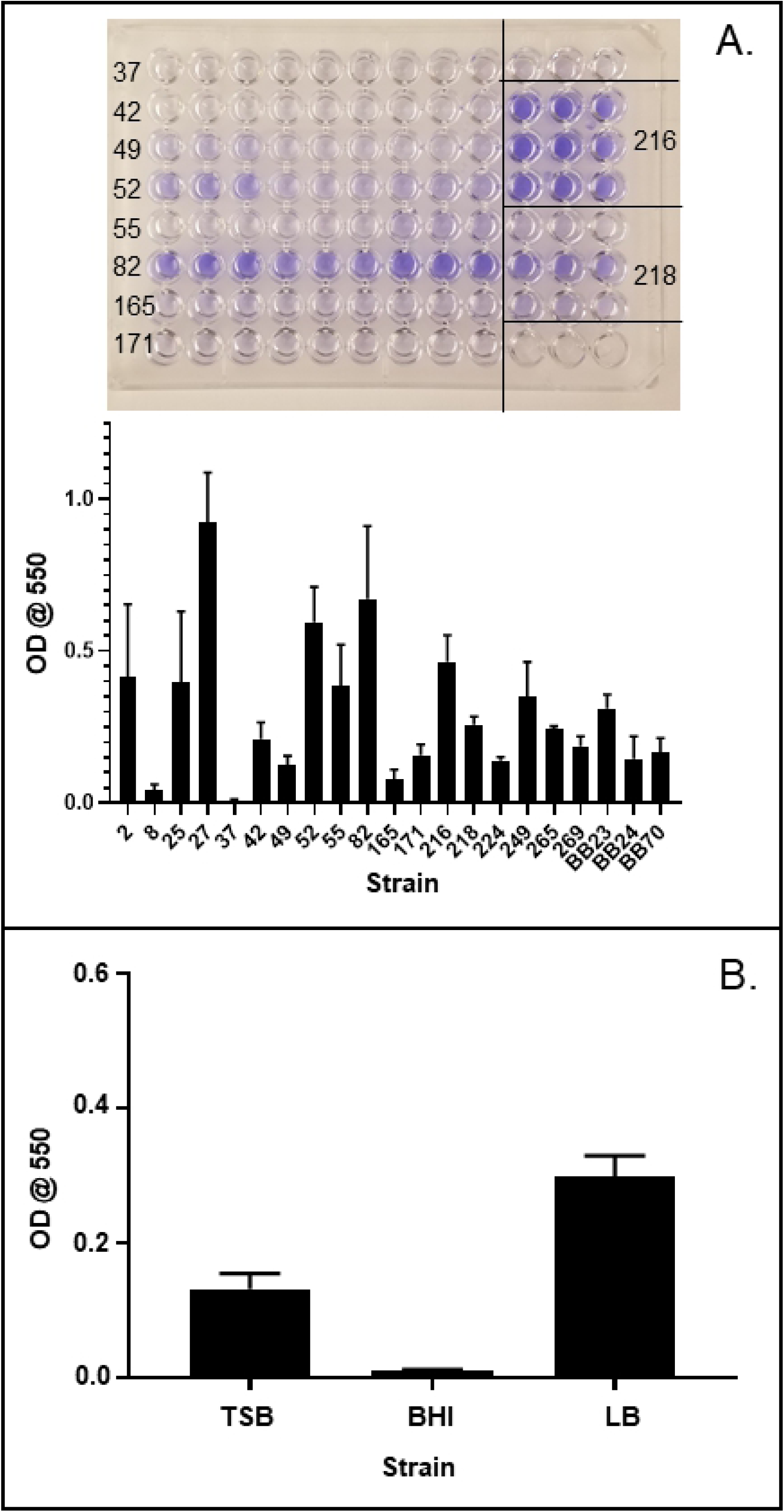
Biofilm formation by *E. faecalis* strains. (A) Representative biofilm assay and average biofilm formation (n=3) for different strains grown in LB. (B) Biofilm formation in strain 82 grown in the indicated media (n=3).

### Maternal enterococci outcompete most artificially introduced strains of *E. faecalis* in colonization of newborn rats

To examine effects of different *E. faecalis* strains in experimental NEC, we introduced these bacteria to newborn rats on day 1 and scored NEC pathology on day 4 of the NEC-inducing regimen of formula feeding-hypoxia. Percentages of *E. faecalis* in intestinal microbiomes and strain composition of *E. faecalis* on day 4 were also determined. Two strains with contrasting combinations of potentially pathogenic phenotypes were chosen for initial experiments. Strain 8 is β-hemolytic, negative for gelatin liquefaction, NF-κB activation, and biofilm formation. Strain 82 is α-hemolytic, positive for gelatin liquefaction, NF-κB activation, and biofilm formation (**S2 Data File**). Newborn rats were given 10^8^ CFU of either strain 8 or strain 82 once, with first formula feed. Control animals were given equivalent amount of bacterial culture supernatant. After 4 d of formula feeding-hypoxia, animals were sacrificed, and intestinal content was plated on blood agar for total bacterial counts and BHI-azide for *E. faecalis. E. faecalis* strains (50-75 azide-resistant colonies per animal) were identified by replica plating onto diagnostic media. NEC was scored microscopically. Interestingly, neither of the two strains was recovered from the inoculated animals; all enterococci isolated were thus of the maternal origin (**S3 Data File**). There were no significant differences in NEC scores between control group and animals inoculated with strains 8 or 82 (**Table 1**). Thus, strains 8 or 82 failed to appreciably colonize neonatal rats upon artificial introduction. Inoculation with these strains did not have significant effect on NEC pathology.

**Table 1.** Distribution of NEC scores in 4-day-old rats

| Treatment group | NEC Score |  |  |  |  | n | p-value* |
| --- | --- | --- | --- | --- | --- | --- | --- |
|  | 0 | 1 | 2 | 3 | 4 |  |  |
| Breast-fed | 7 | 0 | 0 | 0 | 0 | 7 |  |
| Formula-hypoxia (FFH) | 17 | 13 | 11 | 6 | 1 | 48 |  |
| FFH + *E. faecalis* 8 | 29 | 26 | 35 | 9 | 0 | 99 | 0.24 |
| FFH + *E. faecalis* 82 | 9 | 12 | 14 | 6 | 1 | 42 | 0.25 |
| FFH + *E. faecalis* BB70 | 3 | 7 | 10 | 14 | 0 | 34 | <0.0001 |

\*Compared to formula-hypoxia group, $\chi^2$ test.

### Identification of efficient colonizers among *E. faecalis* strains

One reason for the failure of strains 8 and 82 to effectively colonize the neonatal intestine may be adaptive disadvantage of bacteria grown to stationary phase in liquid BHI culture. Indeed, bacteria coming from mothers may successfully colonize the offspring because they are adapted to survival and growth in the organismal environment. In attempts to improve colonization, we tried different culture conditions including growth in medium optimal for biofilm formation (LB), pre-incubation in FBS, or starving bacteria in dilute TSB to induce dormant state. None of these treatments significantly promoted colonization (**S3 Data File**).

In another approach to improving colonization, we set out to determine whether some of our *E. faecalis* strains are inherently better colonizers than others. Newborn rats were given a combined inoculum of all 21 strains mixed in equal proportions, and strain composition of the enterococci was determined on day 4. Strikingly, only 3 strains out of 21 turned out capable of at least some degree of colonization (**Fig 4A**). Strain BB70 was the best colonizer—it was found in all animals that received the mixed inoculum, and constituted, on average, 1/3 of enterococcal populations. None of the input strains were recovered from control non-inoculated animals. We next evaluated efficiency of colonization with pure culture of BB70. In all inoculated animals, *E. faecalis* populations were almost entirely BB70 (**Fig 4B**). Our results indicate that most enterococcal strains failed to colonize newborn rat intestine upon introduction as pure culture. However, some strains could be quite efficient colonizers.

**Figure 4.**
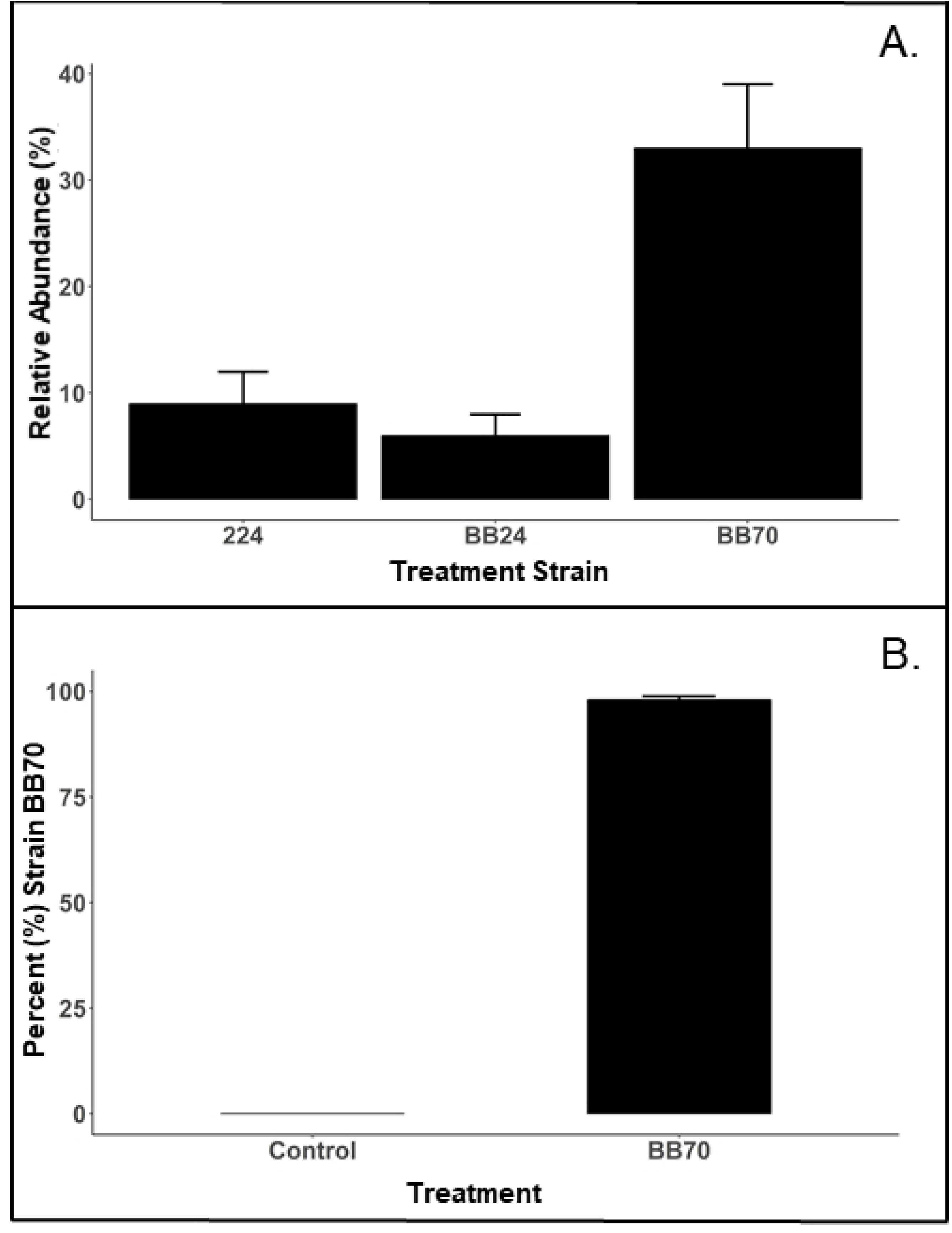
Selection of efficient *E. faecalis* colonizers. (A) Neonatal rats (n = 18) were given oral inoculum containing equal concentrations of all 21 *E. faecalis* strains, followed by 4 d of formula feeding-hypoxia. Only 3 strains (224, BB24, BB70) were recovered on day 4 at indicated average percentages of total *E. faecalis*. (B) Proportion of BB70 in populations of *E. faecalis* in animals that received or did not receive 108 CFU of this strain with first feed (n=21 in each group).

### *E. faecalis* BB70 exacerbates NEC pathology

*E. faecalis* BB70 is negative for hemolysis, gelatin liquefaction, antibiotic resistance, and biofilm, therefore it was expected to be innocuous. However, animals inoculated with this strain had significantly higher NEC scores than control formula-hypoxia animals (**Table 1**), indicating that contrary to expectations, BB70 behaved as a NEC pathogen.

## Discussion

We isolated and characterized 21 different strains of *E. faecalis* from neonatal rats. The strains differed in their colony appearance, hemolysis, gelatin liquefaction, antibiotic resistance, β-galactosidase, and fermentation of sorbitol, mannitol, and arabinose. The strains also differed in their ability to form biofilm and to activate the pro-inflammatory transcription factor NF-κB in cultured enterocytes. There were two genomic variants based on Hin*d*III DNA fragment patterns. Only 3 out of 21 strains appreciably colonized the GI tract of newborn rats upon artificial introduction with first feed. The most efficient colonizer, *E. faecalis* BB70, significantly exacerbated NEC pathology. These results provide an insight into the role of *E. faecalis* in the pathogenesis of experimental NEC.

The strain diversity that we observed was somewhat surprising considering that all animals were from a specific pathogen-free environment at facilities renowned for their high standards of animal care and cleanness. This diversity may indicate that laboratory rat populations harbor a multitude of *E. faecalis* strains with either equal adaptive fitness in the organismal environment, or specific adaptation to different ecological niches. The identification of new strains during the course of our inoculation experiments also suggests that strain composition at the suppliers’ facilities might have changed over the course of several years. Laboratory rats thus present an interesting model to examine significance of the previously described enterococcal diversity [33–35].

*E. faecalis* strains that we isolated originated from the specific pathogen-free environment, therefore none of them is a likely hardcore pathogen. Nevertheless, some of the strains’ phenotypes, such as hemolysis, gelatin liquefaction, antibiotic resistance, biofilm formation, or activation of pro-inflammatory signaling could be associated with opportunistic pathogenicity in the context of NEC. We identified strains possessing multiple potentially pathogenic traits, such as 82, as well as strains with one or no pathogenic traits, such as 8 or BB70. We hypothesized that the former will behave as pathogens and the latter as innocuous or protective symbionts in the rat model of NEC. However, the fact that presumably innocuous BB70 turned in fact pathogenic is contrary to this hypothesis. Unfortunately, we were unable to establish pathogenicity of other strains because of poor colonization. Whether or not potentially pathogenic phenotypes of *E. faecalis* predict increased pathogenicity in vitro remains an open question.

The failure of the majority of our strains to colonize the neonatal intestine upon early introduction was a surprising finding in view of the fact that all the strains were isolated from 4-day-old rats and thus had previous history of successful neonatal colonization. Artificial colonization did not improve significantly by inducing dormancy, culturing in media that promoted biofilm formation, or pre-incubation of bacteria with FBS. A plausible explanation for the poor colonization with bacterial cultures is that maternal enterococci, but not cultured bacteria, are adapted to the organismal environment and therefore have higher probability of survival upon transfer to the neonates. Strain BB70 was an exception: it always outcompeted maternal *E. faecalis* strains. It is possible that in vivo survival of bacteria depends on host-induced genes, and such genes may be constitutively expressed in BB70. Our findings indicate that failure of cultured bacteria to establish intestinal colonization may be a serious limitation to probiotic therapy. Finding probiotic strains similar to BB70 in colonization ability may be a way of overcoming this limitation.

---

**S1 Data File. Phenotypic characterization of 147 isolates of naturally-occurring *E. faecalis***

**S2 Data File. Characteristics of 21 unique strains of *E. faecalis***

**S3 Data File. Bacterial populations and NEC scores of 4-day-old rats following various treatments**

## Acknowledgments

We thank Monica Williams for help in isolating *E. faecalis* strains; Alec Borsook and Alex Li for bacterial binding experiments. This study was supported by NIH grant AI 014032 to H.R.F.

